# TKI Type Switching Overcomes ROS1 L2086F in ROS1 Fusion-Positive Cancers

**DOI:** 10.1101/2024.01.16.575901

**Authors:** Rajat Thawani, Matteo Repetto, Clare Keddy, Katelyn Nicholson, Kristen Jones, Kevin Nusser, Catherine Z. Beach, Guilherme Harada, Alexander Drilon, Monika A. Davare

## Abstract

**Purpose:** Despite the robust efficacy of ROS1 tyrosine kinase inhibitors (TKIs) in ROS1-positive non-small cell lung cancer, TKI resistance continues to hamper durability of the therapeutic response. The resistance liabilities of next-generation ROS1 TKI are sparsely characterized.

**Design:** We compared the activity of type I TKIs (crizotinib, entrectinib, taletrectinib, lorlatinib, and repotrectinib) to the type II TKIs (cabozantinib and merestinib), and to the type I FLT3 inhibitor, gilteritinib, in CD74-ROS1 wildtype and F2004C, L2026M, G2032R, or L2086 mutant Ba/F3 cells. The findings from the Ba/F3 cell model were confirmed using NIH3T3 colony formation assays and in vivo tumor growth. CRISPR/Cas9 gene editing was used to generate isogenic wildtype and L2086F mutant TPM3-ROS1 expressing patient-derived cell lines. These lines were used to further evaluate TKI activity using cell viability and immunoblotting methods. Molecular modeling studies enabled the characterization of structural determinants of TKI sensitivity in wildtype and mutant ROS1 kinase domains. We also report clinical cases of ROS1 TKI resistance that were treated with cabozantinib.

**Results:** ROS1 L2086F mutant kinase is resistant to type I TKI including crizotinib, entrectinib, lorlatinib, repotrectinib, taletrectinib, while the type II TKI cabozantinib and merestinib retain activity. Additionally, we found that gilteritinib, a type I FLT3 inhibitor, inhibited wildtype and L2086F mutant ROS1, however ROS1 G2032R solvent front mutation imposed resistance. The specific binding poses adopted by cabozantinib in the DFG-out kinase conformation and gilteritinib in the DFG-in kinase, provide rationale for their activity in the ROS1 mutants. Clinical cases demonstrated response to cabozantinib in tumors developing TKI resistance due to the ROS1 L2086F mutation.

**Conclusion:** Cabozantinib and gilteritinib effectively inhibit ROS1 L2086F. Clinical activity of cabozantinib is confirmed in patients with TKI-resistant, ROS1 L2086F mutant NSCLC. Gilteritinib may offer an alternative with distinct off-target toxicities, however further studies are required. Since cabozantinib and gilteritinib are multi-kinase inhibitors, there is a persistent unmet need for more selective and better-tolerated TKI to overcome ROS1 L2086F kinase-intrinsic resistance.

**Translational relevance:** ROS1 L2086F is an emerging recurrent resistance mutation to type I ROS1 TKIs, including later generation TKIs. Here, we show corroborating preclinical and clinical evidence for the activity of the quinolone-based type II TKI, cabozantinib, in ROS1^L2086F^ resistance setting. In addition, we show activity of the pyrazine carboxamide-based type I TKI, gilteritinib, in ROS1 L2086F resistance, suggesting that gilteritinib could be another option for ROS1 L2086F TKI-resistant patients. This study represents the first comprehensive report of ROS1 L2086F in the context of later-generation TKIs, including the macrocyclic inhibitors.

## Introduction

Chromosomal rearrangements involving the protooncogene and receptor tyrosine kinase (RTK) *ROS1* generate catalytically active ROS1 fusion oncoproteins^1^. Patients with *ROS1* fusion-containing cancers display exquisite and durable sensitivity to small molecule ROS1 tyrosine kinase inhibitors (TKIs).

TKIs are broadly classified based on their preferred binding mode within the kinase domain. Kinase domains have evolved to adopt a common two-lobe fold (N- and C-l) connected by a hinge. ATP binds within a pocket created between the N- and C-lobes and the conformation of the activation loop (A-loop) which is marked by a conserved Asp-Phe-Gly (“DFG”) motif at its start. In the active conformation, the DFG motif orients towards the ATP-binding site, whereas in the inactive conformation, the DFG motif is flipped to create a catalytically incompetent state unfavorable for ATP binding. Notably, in the DFG-out kinase conformation, a new allosteric binding pocket, adjacent to the ATP binding pocket, is observed. Generally, TKI that occupy the ATP binding pocket are classified as type I or ATP-competitive inhibitors. Among ROS1 TKI, crizotinib, taletrectinib, lorlatinib, repotrectinib and NVL-520 are ATP-competitive type I inhibitors. We previously showed that entrectinib exhibits varying degrees of resistance to ROS1 kinase domain mutations associated with both type I TKI resistance (e.g., ROS1 G2032R) and type II TKI resistance (e.g., ROS1 F2004C), thus suggesting that entrectinib is potentially a type I/II inhibitor with dual binding mode potential^2^. We previously established that the multi-kinase inhibitor, cabozantinib, operates as a type II ROS1 TKI and is liable to resistance with ROS1 D2113N/G, F2004C and F2075C kinase domain mutations^3^. These resistance mutations have not been observed with type I inhibitors, supporting the unique binding pocket requirements for the inhibitors.

In TKI-naïve ROS1 fusion-containing NSCLCs, the three approved ROS1 TKIs (crizotinib, entrectinib, and repotrectinib) achieve high response rates (70-80%) and prolonged progression-free survival (PFS)^4–6^. Next-generation TKIs such as repotrectinib^7^, taletrectinib^8–10^, and NVL-520^11^ were designed to target solvent front resistance arising from the recurrent crizotinib and entrectinib-resistant ROS1 G2032R mutation, with clinical responses to these to these second-line agents now reported in patients with ROS1 G2032R. Furthermore, repotrectinib is now a first line TKI option, achieving a longer median PFS in this setting compared to crizotinib or entrectinib (∼36 months vs ∼16 months for both drugs).

The acquired resistance landscape is likely to adapt to the increasing use of next-generation TKIs. Kinase-intrinsic resistance is typically mediated by the acquisition of *ROS1* mutations with steric, functional, or conformational consequences. We previously reported data from a cohort of patients with *ROS1*-rearranged NSCLC treated with crizotinib and lorlatinib, a National Cancer Center Network Guidelines listed but unapproved ROS1 TKI^12^. The ROS1 G2032R solvent front mutation was most commonly observed. Next-generation TKIs with anti-ROS1 G2032R activity remain type I inhibitors, interacting with the ATP binding site within the kinase’s active pocket like older drugs. As such, resistance mechanisms that broadly affect type I inhibitors may become more clinically relevant.

We and others observed the emergence of on-target resistance due to ROS1 L2086F in *ROS1* fusion-containing cancers treated with both earlier-generation (crizotinib, entrectinib and lorlatinib^12^) and next-generation (taletrectinib) ROS1 TKIs^2,12,13^. L2086F is a catalytic spine 6 (CS6) mutation in ROS1 that appears to disrupt type I TKI binding. As a result, ROS1 L2086F may become a prevalent kinase-intrinsic resistance liability, replacing ROS1 G2032R. Given this potential impact, we sought to characterize ROS1 L2086F and describe therapeutic strategies for this mutation.

## Results

### Activity of ROS1 inhibitors with different binding modes in TKI-resistant ROS1 mutants

Using dose-response cell viability assays in Ba/F3 CD74-ROS1 wild-type and ROS1 F2004C, L2026M, G2032R and L2086F kinase domain mutant cell lines, we determined the cell-based 50% inhibitory concentration (IC_50_) for type I TKI (crizotinib, entrectinib, taletrectinib, lorlatinib, and repotrectinib), and type II TKI (cabozantinib, merestinib), and the type I FLT3 inhibitor, gilteritinib that was recently shown to have activity in ALK fusion driven cancer models^14^ (**Figure 1A**, Supplemental Figure 1). As expected, older ROS1 inhibitors lost potency in cells with the ROS1 G2032R mutation, while next-generation TKIs, taletrectinib and repotrectinib maintained activity. Cell-based IC_50_ for CD74-ROS1 ROS1 L2086F was 50-500-fold higher as compared to CD74-ROS1 ROS1 wildtype in the case crizotinib, entrectinib, lorlatinib, taletrectinib and repotrectinib, establishing this mutation as a resistance liability to all established type I ROS1 TKI. In contrast, this mutation modestly sensitized the protein to cabozantinib, merestinib and gilteritinib; these inhibitors had a lower IC_50_ for ROS1 L2086F than for ROS1 wildtype cells. Intriguingly, ROS1 G2032R was resistant to gilteritinib like the other type I TKIs, confirming that, similar to its binding of FLT3, gilteritinib likely harbors preferential type I binding mode for DFG-in ROS1. Consistent with our previous findings that showed the F2004C mutation imposes resistance to type II TKI like cabozantinib and foretinib, here we confirmed that in addition to cabozantinib and foretinib, the type II TKI merestinib also is resistant to ROS1 F2004C^3^. A network diagram summarizing the kinase-intrinsic resistance liabilities for all the tested TKI, as based on the dose-response cell viability assay data, is shown in **Figure 1B**.

**Figure 1.**
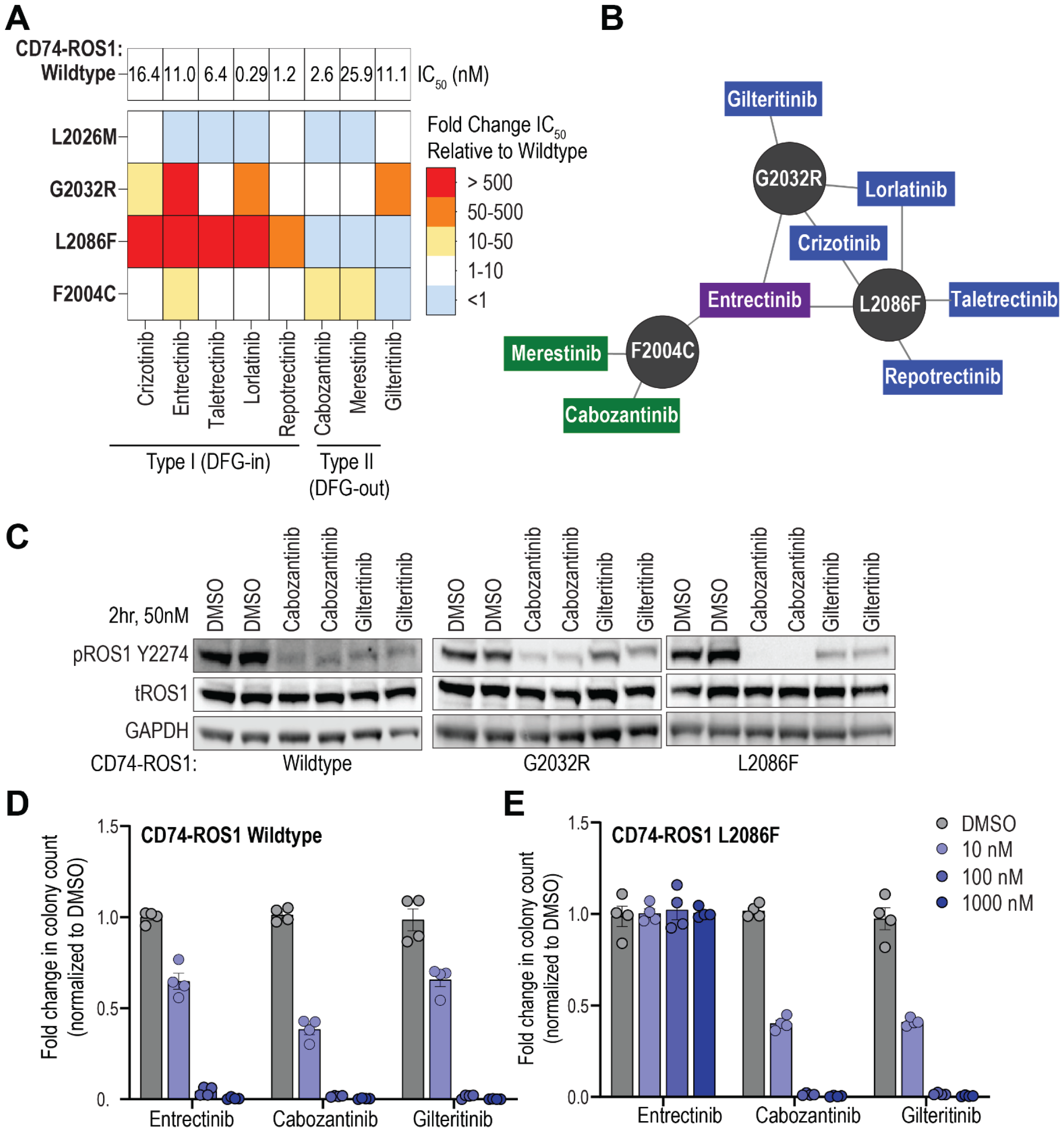
TKI activity in models of ROS1 kinase-intrinsic resistance. **A**. Heat map of IC_50_ values shows relative potency (nM) of the indicated ROS1 inhibitors for Ba/F3 *CD74-ROS1* wildtype and mutants cell lines. IC_50_ values were calculated from three replicates from two independent experiments. **B**. Network diagram summarizes tyrosine kinase inhibitor (TKI) activity against ROS1 G2032R, L2086F and F2004C mutants, with type I inhibitors shown in blue, type II in green and type I/II in purple. **C**. Immunoblot analysis of phosphorylated and total ROS1 in cell lysates generated from Ba/F3 *CD74-ROS1* wildtype, G2032R, and L2086F cell lines treated with indicated inhibitors at 50 nM for 2 hours. **D**. Colony counts anchorage-independent soft agar assays using NIH3T3 *CD74-ROS1* WT and L2086F cell lines. Data normalized to vehicle (DMSO) treatment for indicated TKI treatment conditions. N=4 wells per condition.

Immunoblotting was used to explore the functional consequences of cabozantinib and gilteritinib as they demonstrated robust activity against ROS1 L2086F (**Figure 1C**). Both inhibitors reduced ROS1 autophosphorylation in Ba/F3 CD74-ROS1 wildtype and L2086F mutant cells, correlating with potency in Ba/F3 cell viability assays. Cabozantinib reduced autophosphorylation in ROS1 G2032R, whilst gilteritinib was not effective in case of this mutant, also consistent with the dose response cell viability data.

To confirm the activities of cabozantinib and gilteritinib in an independent model system, we performed soft-agar colony formation assays with NIH3T3 CD74-ROS1 cell lines (WT and L2086F) comparing treatments with entrectinib. Dose dependent inhibition of WT colonies with entrectinib, cabozantinib and gilteritinib was observed (**Figure 1D**). ROS1 L2086F colony formation was resistant to entrectinib but sensitive to cabozantinib and gilteritinib (**Figure 1E**, Supplemental Figure 2A, B). Similar results were observed in cell lines expressing the EZR-ROS1 fusion (Supplemental Figure 2C-F). We tested the in vivo anti-tumor efficacy of cabozantinib and gilteritinib using NIH3T3 CD74-ROS1 WT and L2086F mutant subcutaneous allograft models. Tumor bearing mice were treated with 30 mg/kg cabozantinib or gilteritinib, once daily, and compared to vehicle treatment. This dosing strategy was consistent with previous murine model studies using these agents^14,15^. Cabozantinib and gilteritinib treatments significantly inhibited the growth of both WT ROS1 and ROS1 L2086F tumors (p < 0.05 as indicated, Supplementary Figure 3).

### Activity of non-type I ROS1 TKIs in a patient-derived cell line expressing an endogenous L2086F mutation

Currently, there are no patient-derived *ROS1* fusion-positive cell lines harboring the L2086F resistance mutation. CUTO-28 is a previously establishedTPM3-ROS1-fusion-expressing, patient-derived NSCLC cell line with sensitivity to ROS1 TKIs^16^. We used CRISPR-Cas9 genome editing coupled with homology directed repair (HDR) to generate isogenic ROS1 WT and L2086F mutant CUTO-28 cell lines. The endogenous ROS1 L2086F mutation within TPM3-ROS1, and its expression was confirmed using Sanger sequencing of amplicons generated from gDNA and cDNA harvested from the engineered cells (Supplementary Figure 4).

We compared the efficacy of the panel of inhibitors tested in the Ba/F3 mutant models, crizotinib, entrectinib, lorlatinib, taletrectinib repotrectinib, cabozantinib, merestinib, and gilteritinib in CUTO28 parental (with ROS1 wildtype kinase domain) and CUTO-28 L2086F mutant cells using dose-response cell viability assays (**Figures 2A**, Supplemental Figure 5). These data confirmed resistance of CUTO-28 ROS1 L2086F cells to crizotinib, entrectinib, lorlatinib, taletrectinib and repotrectinib, as compared to parental cells. Cabozantinib, merestinib and gilteritinib retained comparable activity in the wildtype and L2086F mutant cells. **Figure 2B** depicts a 10- to 1000-fold change in the cell-based IC_50_ of the CUTO-28 ROS1 L2086F cells relative to CUTO-28 parental cells. CUTO-28 Control (wildtype ROS1) cells remain unaffected by the mock genome editing process. Immunoblotting shows that a 4-hour treatment with 100 nM inhibitor (crizotinib, repotrectinib, gilteritinib, and cabozantinib) decreases ROS1 autophosphorylation and activation of downstream effectors (SHP2, ERK1/2, STAT3 and S6) in correspondence with the potency observed in dose response cell viability assays (**Figure 2C**). In comparison, robust inhibition of these proteins is observed only after cabozantinib treatment in the CUTO-28 ROS1 L2086F cells, with downstream effector inhibition paralleling the near complete inhibition of ROS1 autophosphorylation (**Figure 2C**). Gilteritinib’s modest activity with 100nM drug dose treatment may reflect the inherently lower potency of gilteritinib to inhibit ROS1 as compared to cabozantinib, given their cell-based growth inhibitor IC_50_s of 33 versus 158 nM for the wildtype kinase, respectively.

**Figure 2.**
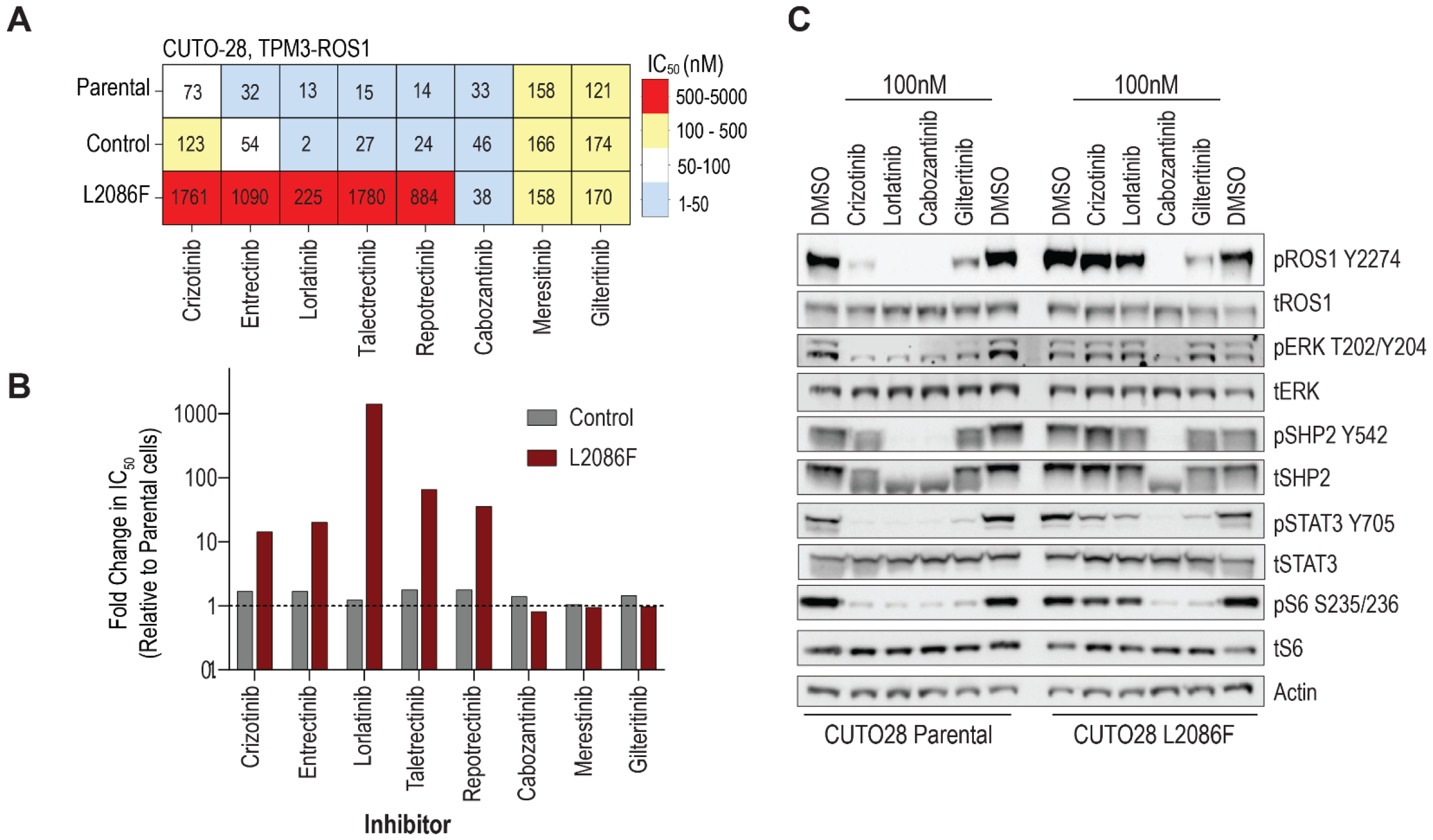
Activity of ROS1 inhibitors in CUTO-28 patient-derived *TPM3-ROS1* fusion cell line with or without L2086F. **A-D**. Cell viability of CUTO-28 *TPM3-ROS1* fusion parental or L2086F cell lines after 72-hour exposure with varying concentrations of crizotinib, cabozantinib, repotrectinib, or gilteritinib as normalized to vehicle-treated cells. Each cell line was plated twice indicated as #1 or #2 except for gilteritinib treatment of the parental CUTO-28 cells. Values are means +/- SEM from 3 replicates. **E**. Immunoblot analysis of phosphorylated and total ROS1, ERK, and downstream pathways in cell lysates generated from CUTO-28 parental and TPM3-ROS1 L2086F cells treated with indicated inhibitors at 100 nM for 4 hours.

### Molecular docking exploration of non-type I TKI interactions with ROS1 wildtype and L2086F mutant kinases

Given the unique pharmacology of gilteritinib compared to type I and type II TKIs, we conducted molecular docking simulation studies to understand the structural basis of the activity of gilteritinib and cabozantinib in ROS1 WT and mutant kinases (**Figure 3**). The studies used the previously reported crystal structure of ROS1 DFG-in kinase (PDB 3ZBF) and homology models of ROS1 DFG-out based on a reported crystal structure of DFG-out ALK (PDB 4FNY). Homology models of ROS1 G2032R and L2086F mutants were also generated.

**Figure 3.**
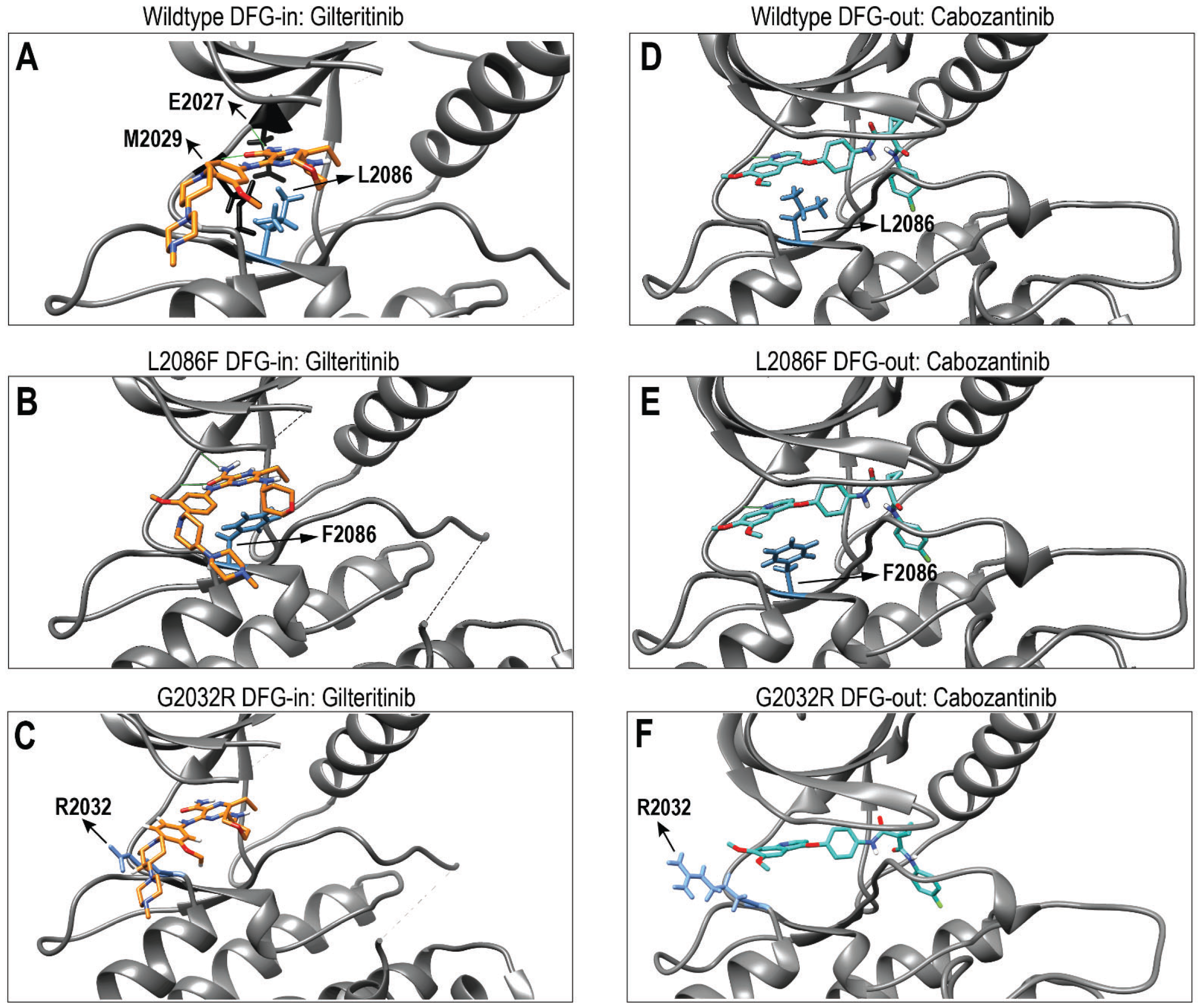
Molecular docking of gilteritinib and cabozantinib to wild-type and mutant ROS1. **A**. Model of wild-type ROS1 kinase domain with gilteritinib in a type I conformation. L2086 is shown in blue and gilteritinib is shown in orange. **B**. Model of wild-type ROS1 kinase domain with cabozantinib bound in a type II conformation. L2086 is shown in blue and cabozantinib is shown in teal. **C**. Model of ROS1 L2086F kinase domain with gilteritinib bound in a type I conformation. F2086 is shown in blue and gilteritinib is shown in orange. **D**. Model of ROS1 L2086F kinase domain with cabozantinib bound in a type II conformation. F2086 is shown in blue and cabozantinib is shown in light teal. **E**. Model of ROS1 G2032R kinase domain with gilteritinib showing a steric clash. (steric clash introduced because of the ROS1 G2032R mutation with gilteritinib bound in the ATP binding pocket of ROS1 is shown in red as predicted in Chimera modeling software.)

The docking of gilteritinib on DFG-in ROS1 was successful for both WT DFG-in ROS1 (**Figure 3A**) and ROS1 L2086F (**Figure 3B**). The model showed that the pyrazine-2-carboxamide moiety of gilteritinib forms two hydrogen bonds with the peptide backbone of both E2027 (H1) and M2029 (H3) in the hinge region of ROS1^17^. This conformation was comparable to those observed in the reported crystal structures of gilteritinib bound to MERTK (PDB 7AB1), and to FLT3 (PDB 6JQR)^18^. This ROS1-giltertinib docked model predicted that gilteritinib occupies the adenosine-binding pocket, the ribose pocket, and the solvent-front cleft of ROS1, confirming it as a type I inhibitor, akin to its reported binding of FLT3. Consistent with cell-based experiments, the bulky R2032 residue from the G2032R solvent-front mutation disrupt optimal gilteritinib binding as it induces steric clash and blocks gilteritinib’s access to the solvent-front cleft (**Figure 3C**), thus making it an ineffective inhibitor for ROS1 G2032R.

As we previously showed, cabozantinib docking was successful on the DFG-out conformation of wildtype ROS1 (**Figure 3D**) and with the ROS1 L2086F (**Figure 3E**) and G2032R (**Figure 3F**) mutants. These models show that the quinoline moiety of cabozantinib forms a hydrogen bond with the H1 residue of ROS1 and extends from the adenosine binding pocket to the back-pocket of the kinase, without substantial extension into the solvent-front cleft.

### Clinical activity of non-type I TKIs in ROS1 L2086F positive NSCLCs

To further investigate the role of drug repurposing to address ROS1 L2086F resistance mutations in the clinical setting, we queried institutional electronic medical records at Memorial Sloan Kettering Cancer Center (MSKCC) to identify patients with acquired ROS1 L2086F mutation following treatment with ROS1 macrocyclic inhibitors. This retrospective query identified four patients, with two of them subsequently enrolled in the prospective phase 2 clinical trial (NCT01639508; IRB 12-097) evaluating cabozantinib efficacy in individuals with advanced NSCLC at MSKCC. This basket trial aims to investigate the role of the type II inhibitor cabozantinib in targeting various acquired resistance mutations based on compelling clinical or preclinical evidence of actionability.

#### Case #1

A 49-year-old otherwise healthy never smoker patient was diagnosed with stage IIIA (ypT1bN2) poorly differentiated lung adenocarcinoma in 2014. The patient received neoadjuvant cisplatin and pemetrexed, followed by right lower lobe lobectomy, and post-operative radiation therapy. In 2015, due to disease recurrence, the patient received first line cisplatin, pemetrexed and bevacizumab with a PR, followed by pemetrexed and bevacizumab maintenance treatment. During chemotherapy, MSK-IMPACT was performed on the surgical specimen and CD74-ROS1 fusion was identified (**Figure 4A**).

**Figure 4.**
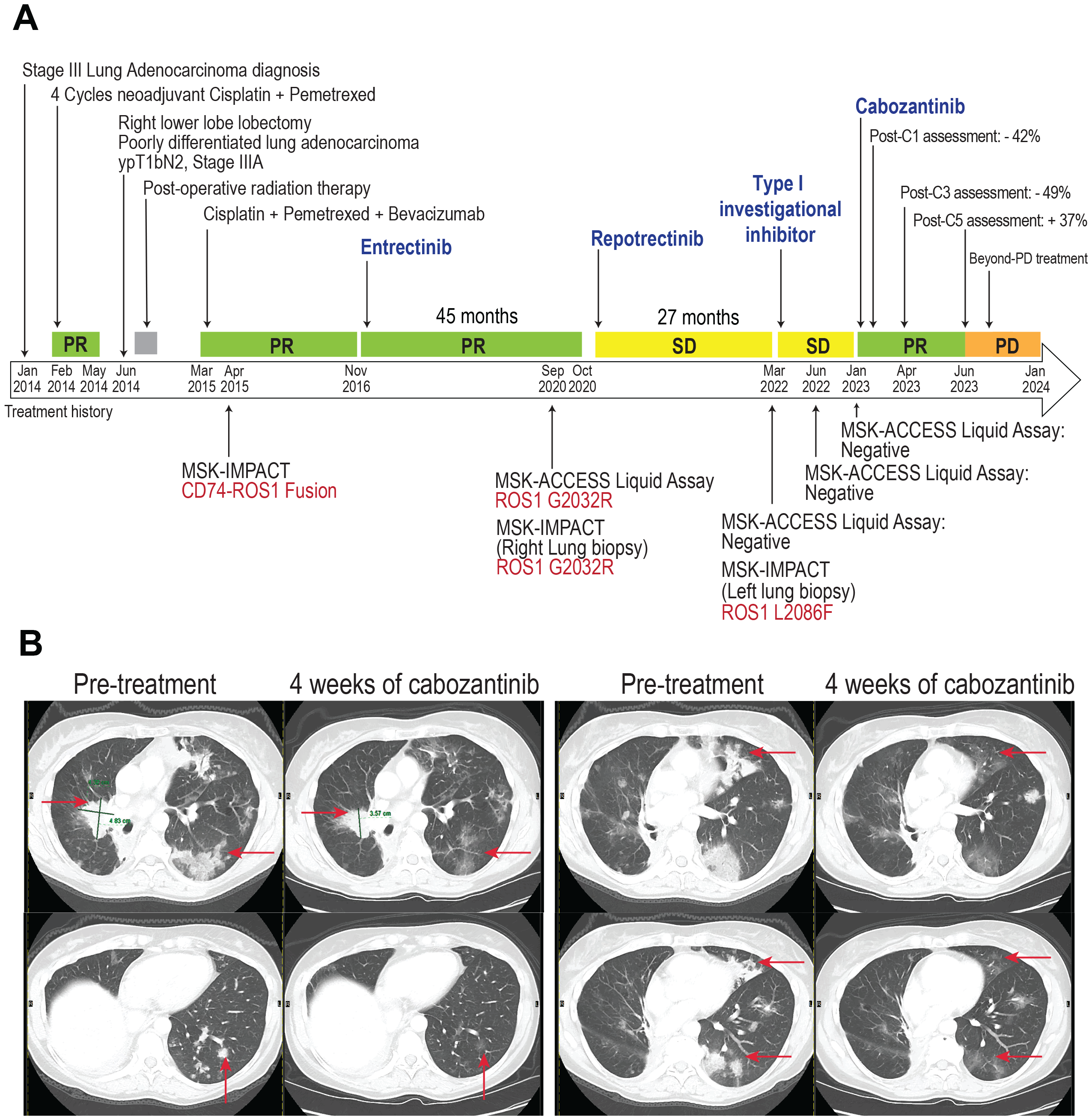
Clinical course of a patient with ROS1 fusion-positive non-small cell lung carcinoma. **A**. Timeline of diagnostic and treatment history of patient which exhibits initial response to cabozantinib. Periods of partial response (PR) are shown in light green, metabolic complete response (CR) shown in green, and stable disease shown in yellow. **B**. Images of chest CT showing partial response to treatment with cabozantinib

The patient then received entrectinib (a type-I inhibitor) from November 2016 to October 2020 with a PR and subsequent PD. Upon PD, both tissue and plasma samples were acquired and analyzed with MSK-IMPACT and MSK-ACCESS (FDA approved plasma ctDNA NGS). Both assays identified an acquired solvent-front ROS1 G2032R mutation. Based on this finding, the patient was subsequently treated with repotrectinib (a type-I macrocyclic inhibitor) for 1 year and 5 months. Upon disease progression on repotrectinib, both tumor and plasma samples were acquired. While circulating tumor (ct)DNA sequencing was negative, tumor sequencing identified loss of the solvent-front ROS1 G2032R mutation and emergence of the CS6 ROS1 L2086F mutation.

The patient was subsequently enrolled in a clinical trial assessing a novel type-I ROS1 selective inhibitor for 10 months. MSK-ACCESS at disease progression in early 2023 was unfortunately negative for detectable somatic alterations. Based on the prior identification of the ROS1 L2086F mutation, the patient was enrolled in a prospective, phase 2 trial of cabozantinib. A RECIST 1.1 partial response (-42% target lesions shrinkage and a dramatic improvement in lymphangitic disease) was achieved after just 4 weeks of therapy, along with a clinical response marked by an improvement in pre-cabozantinib dyspnea. A confirmed partial response was observed on subsequent imaging after 2 months (49% tumor shrinkage, nadir, **Figure 4B**).

Treatment with cabozantinib required a dose reduction from 60 mg daily to 40 mg daily for asthenia, and no major tolerability issues were identified thereafter. After 4 months of therapy, asymptomatic radiological evidence of disease progression was observed with slowly growing pulmonary disease. Due to the overall modest increase in tumor burden, and the enduring clinical benefit reported by the patient, the patient was kept on cabozantinib therapy and remains on the clinical trial more than 12 months into treatment.

#### Case #2

A 70-year-old patient without a smoking history was diagnosed with stage IV NSCLC in 2016 and received first line carboplatin and pemetrexed treatment from September 2016 to April 2017. Following the identification of CD74-ROS1 fusion, the patient was switched to crizotinib (type I inhibitor) and had a partial response (PR) for almost 3 years with subsequent progressive disease (PD). Following progression, the patient was treated with repotrectinib (type I macrocyclic inhibitor) and responded for 1 year and 2 months, however, PD was later observed.

A right lower lobe biopsy of a progressing lesion was collected and analyzed with MSK-IMPACT, an FDA approved next-generation sequencing assay designed for fixed paraffin embedded samples. This resulted in identification of an acquired ROS1 L2086F resistance mutation. The patient was then received cabozantinib (a type II inhibitor) on the same phase 2 trial. The patient’s previously progressive disease stabilized with cabozantinib [best response of stable disease (SD) per RECIST1.1], with a total time on therapy of almost 5 months. The treatment was moderately well tolerated but was complicated by renal infarction which required investigational treatment interruption and subsequent dose reduction. Cabozantinib was ultimately discontinued due to RECIST1.1 PD; the patient did not receive additional treatments.

## Discussion

The study of kinase-intrinsic resistance has been crucial to improving outcomes in patients with advanced, fusion-driven cancers. Joining *BCR-ABL-*positive chronic myelogenous leukemia and *ALK* fusion-positive NSCLCs, *ROS1* fusion-positive NSCLCs are a recent example of how sequential TKI therapy (i.e., using repotrectinib in TKI pre-treated patients) can extend the total duration of disease control for an individual. Notably, the patterns of on-target or kinase intrinsic resistance tend to reflect the TKI-binding mode driven selection pressure, wherein cases treated with type I inhibitors showed prevalence of the solvent front, ROS1 G2032R mutation to date. On target resistance is driven by expansion of tumor cells harboring TKI type-specific resistant mutations. While repotrectinib, with activity on ROS1 G2032R resistance, is the only approved second line TKI option, other inhibitors such as taletrectinib and NVL-520 are also highly active and may be approved in the future. The increasing use of next-generation TKI therapy is likely to shift emergent resistance patterns.

In this study, we present preclinical and clinical data supporting the hypothesis that ROS1 L2086F is a convergent liability for type I ROS1 TKIs; these include approved (crizotinib, entrectinib, repotrectinib), guidelines-listed (lorlatinib), and investigational (taletrectinib) agents. Most importantly, we provide information on a therapeutic strategy for ROS1 L2086F using agents such as gilteritinib (for which this series provides the first data supporting use) and cabozantinib, both of which are available in the clinic and approved for other indications. As a classical type II inhibitor, cabozantinib binding is unaffected by solvent front mutations such as ROS1 G2032R and the recalcitrant central-sheet 6 mutation, ROS1 L2086F. Our data showed that gilteritinib exhibited unexpected behavior as a type I TKI such that ROS1 G2032R imposes resistance to this inhibitor, however, owing to its specific binding pocket occupancy within the ROS1 kinase domain, it remains unhindered by ROS1 L2086F. The use of one over the other would be informed by a variety of factors, including side effects (the drugs have vastly different toxicity profiles) and kinase-intrinsic resistance concurrent with ROS1 L2086F. Specifically, cabozantinib may retain activity in the presence of a ROS1 G2032R while gilteritinib is not predicted to be active.

The overarching observation is that TKI type switching is a legitimate method of abrogating select pan type I TKI kinase-intrinsic resistance in ROS1 fusion driven NSCLC. In contrast to type I TKIs, type II TKIs bind a hydrophobic pocket adjacent to the ATP-binding region when the kinase activation loop (A-loop) adopts the “DFG-out” conformation, thus bypassing the solvent front mutations that pose challenges for type I inhibitors. We and others previously demonstrated the benefits of type II TKI use with ROS1 D2033N resistance, and in *NTRK* fusion-positive cancers that acquire xDFG (TRKA G667, TRKC G696) resistance substitutions that stabilize the DFG-out conformation. Gilteritinib was active against the ALK L1256F resistance mutation in *ALK* fusion-positive NSCLCs, and against select D1228 resistance substitutions in MET-driven cancers that progressed on prior type I TKI therapy^19^. Hence, sequencing-informed TKI type switching has the potential to improve outcomes and provide effective therapeutic interventions in many resistance-driven cancer settings^20^.

Unfortunately, in fusion-positive cancers, few rationally designed next-generation type II inhibitors have been developed. All approved agents for any fusion-positive cancer (e.g., involving *ALK, RET, ROS1, NTRK1/2/3, FGFR1/2/3*) are type I drugs. The successful development of novel type II inhibitors may thus provide a transformative opportunity for targeting kinase-intrinsic resistance.

## Materials and Methods

### Cell Culture

Ba/F3 and NIH3T3 cells were purchased from American Type Culture Collection. CUTO-28 cells were obtained as part of a research agreement from the University of Colorado, following Institutional Review Board approved informed consent of the patient, and has been described by McCoach et al^21^. CUTO-28 cells were cultured in RPMI medium 1640 with 10% (vol/vol) FBS, 2mmol/L L-glutamine, penicillin, streptomycin, and amphotericin B. Parental Ba/F3 cells were cultured in RPMI medium 1640 with 10% (vol/vol) FBS, 2mmol/L L-glutamine, penicillin, streptomycin, amphotericin B and 2ng/mL recombinant murine IL-3. NIH3T3 cells were cultured in DMEM medium with 10% (vol/vol) CS, 2mmol/L L-glutamine, penicillin, streptomycin, and amphotericin B. All cells were tested for mycoplasma at least every three months by using the Lonza MycoAlert™ PLUS Kit.

### Ba/F3 and NIH3T3 cell lines

Generation of Ba/F3 stable cell lines has been described^2^. Cell lines were maintained at densities of 0.5 × 10^6^ to 1.5 × 10^6^ /mL and infected with retrovirus-encoding human *CD74-ROS1* and *EZR-ROS1* fusion genes. *CD74-ROS1* and *EZR-ROS1* fusions with F2004C, L2026M, G2032R, L2086F mutations were made using site-directed mutagenesis following the manufacturer’s protocol (Agilent) using primers listed in Supplementary Table S1. Platinum-E cells (Cell Biolabs, Inc) were transfected with pBABE *CD74-ROS1* or pCX4 *EZR-ROS1* wild-type or mutant constructs using Biotool DNA transfection reagent to generate replication incompetent, ecotropic retrovirus. Parental NIH3T3 cells were seeded at 50% confluence and infected with retrovirus. Cells were selected with puromycin treatment (2 μg/ml) for 72-96 hours prior to being used for experimental work. Transformed Ba/F3 cell lines: For IL-3 withdrawal, the cells were washed three times with complete medium without IL-3, and seeded at 0.5 x 10^6^ cells per mL in RPMI medium 1640 with 10% (vol/vol) FBS, 2mmol/L L-glutamine, penicillin, streptomycin, amphotericin B without supplemented IL-3. Cells were counted every 2 days until the population had at least tripled indicating that the proliferating population was IL-3 independent. Sanger sequencing of integrated cDNA in the transformed cell lines was done to verify the presence of the desired mutation. For this, DNA was harvested from cell pellets using QuickExtract DNA Extraction Solution (Lucigen). The ROS1 kinase and C-terminal domains were PCR amplified using the primers ROS1 5707F (5’-GACAAAGAGTTGGCTGAGCTG-3’) and ROS1 REV_14 (5’-TCAGACCCATCTCCATATCCA-3’) and bidirectionally sequenced using the primers ROS1 6198F (5’-CTGTGTCTACTTGGAACGGATG-3’) and ROS1 6304R (5’-TCTCTGGCGAGTCCAAAGTC-3’). Chromatographs were aligned using Benchling software to verify that the correct mutation was present, and no other mutations were introduced during viral transduction.

### Generation of mutated patient-derived cell line

The CUTO-28 cell line is a patient-derived cell line harboring a *TPM3-ROS1* fusion, authenticated by short tandem repeat (STR) analysis by the Barbara Davis Center Molecular Biology Service Center at the University of Colorado^16^.

A CRISPR-Cas9 genome-engineered CUTO28 cell line with an endogenous L2086F mutation was engineered. A pX330 vector was made to express a guide directed to the L2086 region of ROS1. A homology directed repair (HDR) oligo template was designed to have 80 base pairs of homology on each side of the Cas9 cut site with a single base pair substitution allowing for the desired mutation and alteration of the PAM site. CUTO28 cells were plated at a density of 500,000 cells per well, in 6 well plate, 24 hours prior to transfection, then transfected with either GFP alone or 1 ug pX330(I) L2086F guide and 10 ug HDR Oligo. Forty-eight hours later the media was removed and replaced with RPMI1640 10% Fetal Bovine Serum 1% penstrep 1% L glutamine (R10) containing 500nM lorlatinib and 500nM crizotinib and allowed to grow out. Media with these inhibitors was changed every 72-96 hours. On day 16, cells were placed in inhibitor-free media and allowed to grow in R10. Lucigen QuickExtract reagent was used to generate genomic DNA, and RNA was generated using the Qiagen RNeasy kit, followed by cDNA synthesis using Applied Biological Materials (ABM) All in one 5X Master Mix. Genomic DNA and cDNA were used in polymerase chain reaction to amplify region of interest containing the L2086 codon. Sanger sequencing of the PCR amplicons were used to confirm the genomic edit leading to achieve the desired codon change. Guide DNA, HDR, and PCR primer sequences are reported in Supplemental Table S1.

### Cell Viability Assays

All inhibitors were prepared as 1 mM stocks in dimethyl sulfoxide (DMSO). Inhibitors were distributed at 2X indicated final concentrations into 384-well plates pre-seeded with 25 μl per well of complete medium using a D300 Digital Dispenser (Hewlett-Packard). Ba/F3 cells expressing wild-type or mutant *CD74-ROS1* constructs were seeded at 1,000 cells per well in a volume of 25 μl using a Multidrop Combi Reagent Dispenser (Thermo Fisher Scientific). Plates were incubated for 72 hours at 37°C, 5% CO_2_. Viability was measured using the cell counting kit-8 (CCK-8, Bimake) and read on a Biotek Synergy 2 plate reader. Each condition was assayed in triplicate. Data were normalized using Microsoft Excel, and IC_50_ values were calculated using a nonlinear regression analysis in GraphPad Prism.

### Colony formation assays

Plates were pre-seeded with 0.4% agarose in complete medium [DMEM with 10% (vol/vol) calf serum, 2mM L-glutamine, penicillin, streptomycin, and amphotericin B] with indicated concentrations of each inhibitor. Each inhibitor had its own paired DMSO (vehicle) control condition. NIH3T3 cells expressing *CD74-ROS1* or *EZR-ROS1* wild-type or L2086F were plated in 0.2% agarose in complete medium with indicated concentrations of each inhibitor at a density of 8000 cells per 0.5mLs of agarose; 24 hours after plating 1mL of complete medium with indicated concentrations of each inhibitor was added to each well to prevent dehydration of the agarose. Plates were read 4 weeks after seeding using a GelCount (Oxford Optronix). Colony counts were determined for each well using the GelCount software and normalized to the average colony count from the paired DMSO condition for each inhibitor. Data analysis and visualization was performed using Microsoft Excel and GraphPad Prism.

### Immunoblot analysis

Ba/F3 and CUTO-28 cells were treated with the indicated concentrations of inhibitors for indicated duration, prior to harvest. Ba/F3 cell lines and CUTO28 cells were pelleted, washed with ice-cold phosphate-buffered saline (PBS), and lysed in cell lysis buffer supplemented with 0.25% deoxycholate, 0.05% SDS, and protease and phosphatase inhibitors. Protein concentration was determined using the Pierce BCA Protein Assay Kit (Thermo Fisher Scientific). After protein quantification, lysates were extracted with Laemelli sample buffer for 10 minutes at 75°C and lysates were run on 4% to 12% Bis-Tris gels (Invitrogen; Thermo Fisher Scientific). Proteins were transferred to nitrocellulose membranes (Prometheus) and probed with phospho-ROS1 Y2274 (3078; 1:1,000; Cell Signaling Technology), ROS1 (D4D6; 1:1,000; Cell Signaling Technology), phospho-p44/42 MAPK (9101; 1:1,000, Cell Signaling Technology), ERK2 (sc-1647; 1:1000; Santa Cruz Biotechnology), GAPDH (OTI2D9; 1:5,000; Origene), and Actin (JLA20; 1:5,000; Developmental Studies Hybridoma Bank). Signal was detected using a BioRad ChemiDoc imaging station or a LI-COR Odyssey imaging system with use of HRP-conjugated or IR dye secondary antibodies, respectively.

### Structural modeling

The chemical structures of gilteritinib and cabozantinib were downloaded from PubChem to be employed in docking simulations. Protein complexes 6JQR and 7AB1, which feature Gilteritinib in complex with FLT3 and MerTK kinases respectively, were retrieved from the Protein Data Bank (PDB) to serve as structural references during the docking result analysis phase. Due to the absence of cabozantinib complexes in PDB, the 6SD9 complex of the closely-related quinoline-based type II inhibitor foretinib with MET was obtained from PDB to be used as the closest available reference.

For the target ROS1 kinase, the active conformation (DFG-in) was obtained from PDB (3ZBF), while its inactive state was acquired from previous structural modeling experiments where it was built using ALK (PDB entry 4FNY) DFG-out as a structural template.^22^ The ROS1 L2086F and G2032R mutations were introduced both on the active DFG-in and on the inactive DFG-out ROS1 structures through single amino-acid substitution using PyMOL version 2.5.3. Subsequently, target protein structures, were prepared using Schrödinger Suite and any missing loops or hydrogen atoms were added.

Docking was performed with Glide in Schrödinger Maestro Version 12.7.161. A 30-angstrom grid box was centered around both the WT L2086 and the mutant L2086F CS6 residue. Based on the analysis of gilteritinib and foretinib reference binding poses hydrogen bonding constraints were placed. For gilteritinib, the generation of at least 1 hydrogen bond with either hinge 1 (H1) E2027 or hinge 3 (H3) M2029 peptide backbone was placed, while for cabozantinib generation of a hydrogen bond with H1 residue was required.

Ligands were prepared using the Ligprep and docking simulations were performed using the Glide module in standard precision mode, with the pre-specified hydrogen bonding constraints. Successful binding poses were analyzed and the best scoring ones were selected.^23^

### Murine Model Studies

All animal model studies were conducted in accordance with the Animal Welfare Act (AWA), Public Health Service (PHS), the United States Department of Agriculture (USDA), and under auspices of an approved protocol from the OHSU Institutional Animal Care and Use Committee (IACUC). Four-to six-week-old female athymic nude mice (Nu/J, Strain # 002019, RRID:IMSR_JAX:002019) were purchased from The Jackson Laboratory (Bar Harbor, ME) and housed and handled under specific pathogen-free conditions in the Oregon Health & Science University’s animal care facilities. After an initial 2-week environmental adjustment period, mice were placed under anesthesia using 2% isoflurane/oxygen, weighed, and ear punched for identification purposes. Tumor cells (1 x 10^6^ were mixed with 50 μl of matrigel and injected subcutaneously into the left or right flank. Animals were allowed to recover under supervision before being returned to animal facilities. Injected animals were checked daily until tumors were palpable nodules, at which time both animal weight and tumor size were measured thrice weekly using balance and a digital caliper (cat 14-648-17, Fisher Scientific, Federal Way, WA). Tumor volume was estimated using caliper measurements, based on the ellipsoid volume formula (π/6 x D3). TKI treatment began ∼12-14 days after initial injection when animals were randomized to TKI treatment groups. The study was not blinded. Both gilteritinib and cabozantinib treatment groups were administered indicated TKI at 30 mg/kg formulated in 0.5% methylcellulose, once daily via oral gavage for ten days. Vehicle (0.5% methylcellulose) treated tumors were allowed to grow until they reached the humane limit of 1,500–2,000 mm^3^ at which time animals were sacrificed in accordance with IACUC protocol.

### Patient Clinical Cases

The patients described in this analysis were treated in the prospective phase 2 clinical trial NCT01639508 (IRB 12-097) and provided written informed consent for all procedures described.

### Statistical Analysis

In all graphs, mean ± standard error of means is shown unless otherwise stated with a minimum of three technical replicates. A two-way ANOVA with multiple comparisons tests were used to assess significant differences in tumor volumes in the vivo study. P values < 0.05 were deemed statistically significant. All data were initially analyzed using Microsoft Excel. GraphPad Prism v9.3 software was used for statistical analysis and plotting graphs.

## Acknowledgements

The study was funded by the Medical Research Foundation of Oregon Early Career Investigator Award (RT), NIH/NCI R01CA233495 (Davare), NIH/NCI P30CA008748 (Repetto, Harada, Drilon), NIH/NCI R01CA273224 (Drilon).

## Competing interests

Dr. Drilon reports Honoraria/Advisory Boards: 14ner/Elevation Oncology, Amgen, Abbvie, ArcherDX, AstraZeneca, Beigene, BergenBio, Blueprint Medicines, Chugai Pharmaceutical, EcoR1, EMD Serono, Entos, Exelixis, Helsinn, Hengrui Therapeutics, Ignyta/Genentech/Roche, Janssen, Loxo/Bayer/Lilly, Merus, Monopteros, MonteRosa, Novartis, Nuvalent, Pfizer, Prelude, Repare RX, Takeda/Ariad/Millenium, Treeline Bio, TP Therapeutics, Tyra Biosciences, Verastem; Associated Research to Institution: Foundation Medicine, Teva, Taiho, GlaxSmithKlein; Equity: mBrace, Treeline; Copyright: Selpercatinib-Osimertinib (pending); Royalties: Wolters Kluwer, UpToDate, Food/Beverage: Boehringer Ingelheim, Merck, Puma: CME Honoraria: Answers in CME, Applied Pharmaceutical Science, Inc, AXIS, Clinical Care Options, EPG Health, Harborside Nexus, I3 Health, Imedex, Liberum, Medendi, Medscape, Med Learning, MJH Life Sciences, MORE Health, Ology, OncLive, Paradigm, Peerview Institute, PeerVoice, Physicians Education Resources, Remedica Ltd, Research to Practice, RV More, Targeted Oncology, TouchIME, WebMD. Dr. Davare reports consulting for Nuvalent. The remaining authors report no conflicts of interest.

## References

1. Yu, Z.-Q. et al. ROS1-positive non-small cell lung cancer (NSCLC): biology, diagnostics, therapeutics and resistance. J. Drug Target. 30, 845–857 (2022).

2. Keddy, C. et al. Resistance Profile and Structural Modeling of Next-Generation ROS1 Tyrosine Kinase Inhibitors. Mol. Cancer Ther. 21, 336–346 (2022).

3. Davare, M. A. et al. Structural insight into selectivity and resistance profiles of ROS1 tyrosine kinase inhibitors. Proc. Natl. Acad. Sci. 112, E5381–E5390 (2015).

4. Mazieres, J. et al. Immune checkpoint inhibitors for patients with advanced lung cancer and oncogenic driver alterations: results from the IMMUNOTARGET registry. Ann. Oncol. 30, 1321–1328 (2019).

5. Shaw, A. T. et al. Crizotinib in ROS1-rearranged advanced non-small-cell lung cancer (NSCLC): updated results, including overall survival, from PROFILE 1001. Ann. Oncol. Off. J. Eur. Soc. Med. Oncol. 30, 1121–1126 (2019).

6. Drilon, A. et al. Entrectinib in ROS1 fusion-positive non-small-cell lung cancer: integrated analysis of three phase 1–2 trials. Lancet Oncol. 21, 261–270 (2020).

7. Drilon, A. et al. Repotrectinib (TPX-0005) Is a Next-Generation ROS1/TRK/ALK Inhibitor That Potently Inhibits ROS1/TRK/ALK Solvent-Front Mutations. Cancer Discov. 8, 1227–1236 (2018).

8. Nagasaka, M. et al. TRUST-II: a global phase II study of taletrectinib in ROS1-positive non-small-cell lung cancer and other solid tumors. Future Oncol. 19, 123–135 (2023).

9. Katayama, R. et al. The new-generation selective ROS1/NTRK inhibitor DS-6051b overcomes crizotinib resistant ROS1-G2032R mutation in preclinical models. Nat. Commun. 10, 3604 (2019).

10. Papadopoulos, K. P. et al. U.S. Phase I First-in-human Study of Taletrectinib (DS-6051b/AB-106), a ROS1/TRK Inhibitor, in Patients with Advanced Solid Tumors. Clin. Cancer Res. 26, 4785–4794 (2020).

11. Drilon, A. et al. NVL-520 Is a Selective, TRK-Sparing, and Brain-Penetrant Inhibitor of ROS1 Fusions and Secondary Resistance Mutations. Cancer Discov. 13, 598–615 (2023).

12. Lin, J. J. et al. Spectrum of Mechanisms of Resistance to Crizotinib and Lorlatinib in ROS1 Fusion– Positive Lung Cancer. Clin. Cancer Res. 27, 2899–2909 (2021).

13. Drilon, A. et al. ROS1-dependent cancers — biology, diagnostics and therapeutics. Nat. Rev. Clin. Oncol. 18, 35–55 (2021).

14. Mizuta, H. et al. Gilteritinib overcomes lorlatinib resistance in ALK-rearranged cancer. Nat. Commun. 12, 1261 (2021).

15. Yakes, F. M. et al. Cabozantinib (XL184), a Novel MET and VEGFR2 Inhibitor, Simultaneously Suppresses Metastasis, Angiogenesis, and Tumor Growth. Mol. Cancer Ther. 10, 2298–2308 (2011).

16. Tyler, L. C. et al. MET gene amplification is a mechanism of resistance to entrectinib in ROS1+ NSCLC. Thorac. Cancer 13, 3032–3041 (2022).

17. Pflug, A. et al. A-loop interactions in Mer tyrosine kinase give rise to inhibitors with two-step mechanism and long residence time of binding. Biochem. J. 477, 4443–4452 (2020).

18. Kawase, T. et al. Effect of Fms-like tyrosine kinase 3 (FLT3) ligand (FL) on antitumor activity of gilteritinib, a FLT3 inhibitor, in mice xenografted with FL-overexpressing cells. Oncotarget 10, 6111–6123 (2019).

19. Mizuta, H. et al. Gilteritinib overcomes lorlatinib resistance in ALK-rearranged cancer. Nat. Commun. 12, 1261 (2021).

20. Fujino, T. et al. Foretinib can overcome common on-target resistance mutations after capmatinib/tepotinib treatment in NSCLCs with MET exon 14 skipping mutation. J. Hematol. Oncol.J Hematol Oncol 15, 79 (2022).

21. McCoach, C. E. et al. Resistance Mechanisms to Targeted Therapies in ROS1+ and ALK+ Non–small Cell Lung Cancer. Clin. Cancer Res. 24, 3334–3347 (2018).

22. Drilon, A. et al. A Novel Crizotinib-Resistant Solvent-Front Mutation Responsive to Cabozantinib Therapy in a Patient with ROS1-Rearranged Lung Cancer. Clin. Cancer Res. Off. J. Am. Assoc. Cancer Res. 22, 2351–2358 (2016).

23. Repasky, M. P., Shelley, M. & Friesner, R. A. Flexible ligand docking with Glide. Curr. Protoc. Bioinforma. Chapter 8, Unit 8.12 (2007).

