## Supplementary Figures and Table for "TKI Type Switching Overcomes ROS1 L2086F in ROS1 Fusion-Positive Cancers"

**Supplemental Figure 1**. **Dose response curves characterizing tyrosine kinase inhibitor sensitivity of ROS1 wild-type (WT) and mutant cell lines.** Cell viability of Ba/F3 *CD74-ROS1* WT and mutant cells lines after 72-hour exposure with varying concentrations of crizotinib, entrectinib, taletrectinib, lorlatinib, repotrectinib, cabozantinib, merestinib, and gilteritinib as normalized to vehicle-treated cells. Graphs showing mean +/- SEM from three replicates are representative data from two independent experiments.

**Supplemental Figure 2**. **Soft agar colony formation assays using NIH3T3 *EZR-ROS1* wild-type (WT) and L2086F cells. A-D.** Representative images from colony formation soft agar assays using NIH3T3 CD74::ROS1 and EZR::ROS1 WT and L2086F cells in indicated concentrations of inhibitor. **E-F.** Colony count normalized to paired dimethyl sulfoxide (DMSO) conditions for indicated tyrosine kinase inhibitor treatment conditions from anchorage-independent soft agar assays using NIH3T3 EZR::ROS1 WT or L2086F cell lines. Data from n=4 wells per condition.

**Supplemental Figure 3. In vivo antitumor efficacy of cabozantinib and gilteritinib.** Time course of tumor growth of allografted NIH3T3 *CD74-ROS1* wild-type (A) and 2086F (B) subcutaneous tumors treated with cabozantinib and gilteritinib. N = 4 mice per treatment group. Two-way ANOVA analysis was used to determine significant differences.

**Supplemental Figure 4. Confirmation of L2086F alteration in CUTO-28 cells.**
Sanger sequencing from amplicons generated from CUTO-28 L2086F cell line gDNA (**A)** and cDNA (**B**) confirm the heterozygous L2086F mutation in this gene edited cell line.

**Supplemental Figure 5**. **Dose response curves characterizing tyrosine kinase inhibitor sensitivity of CUTO-28 Parental, CUTO-28 Control HDR edited and CUTO-28 L2086F mutant cell lines.** Cell viability of indicated cells lines after 72-hour exposure with varying concentrations of crizotinib, entrectinib, taletrectinib, lorlatinib, repotrectinib, cabozantinib, merestinib, and gilteritinib as normalized to vehicle-treated cells. Graphs showing mean +/- SEM from three replicates are representative data from two independent experiments.

**Supplementary Table S1.** Primer sequences for site directed mutagenesis, subcloning and PCR amplification reported in the manuscript.

### Supplemental Figure 1

#### Ba/F3 CD74-ROS1

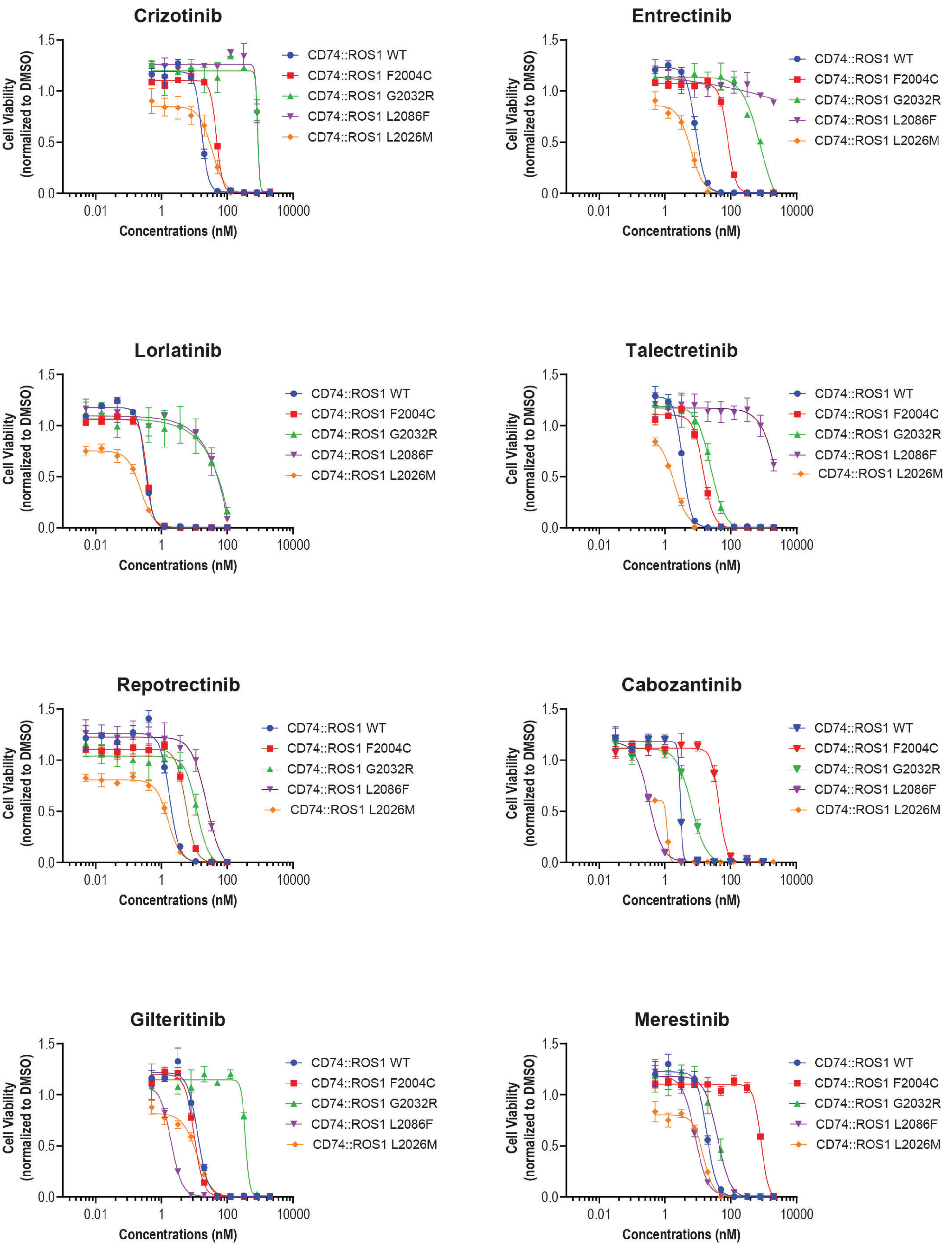

#### Supplemental Figure 2

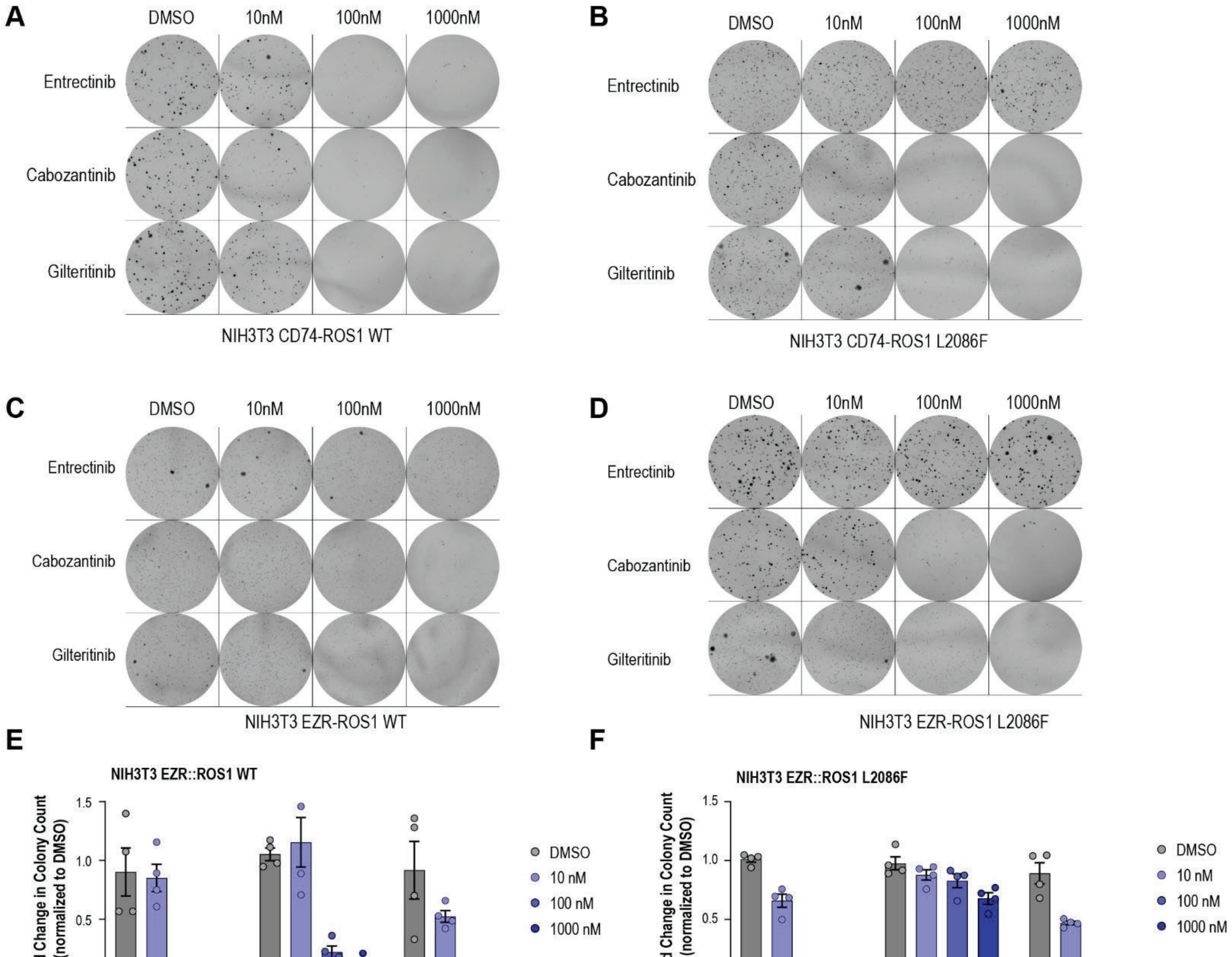

**Supplementary Figure 3**

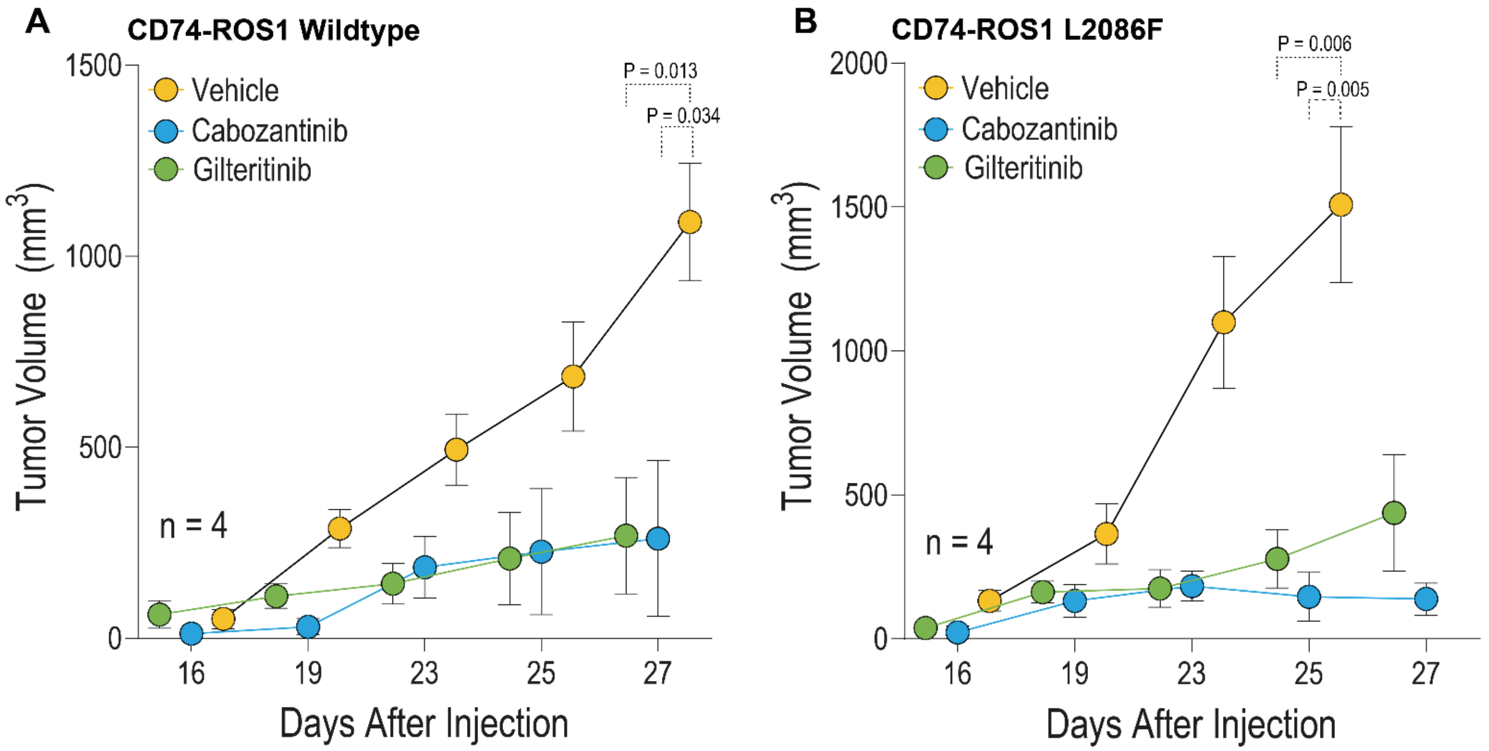

### Supplementary Figure 4

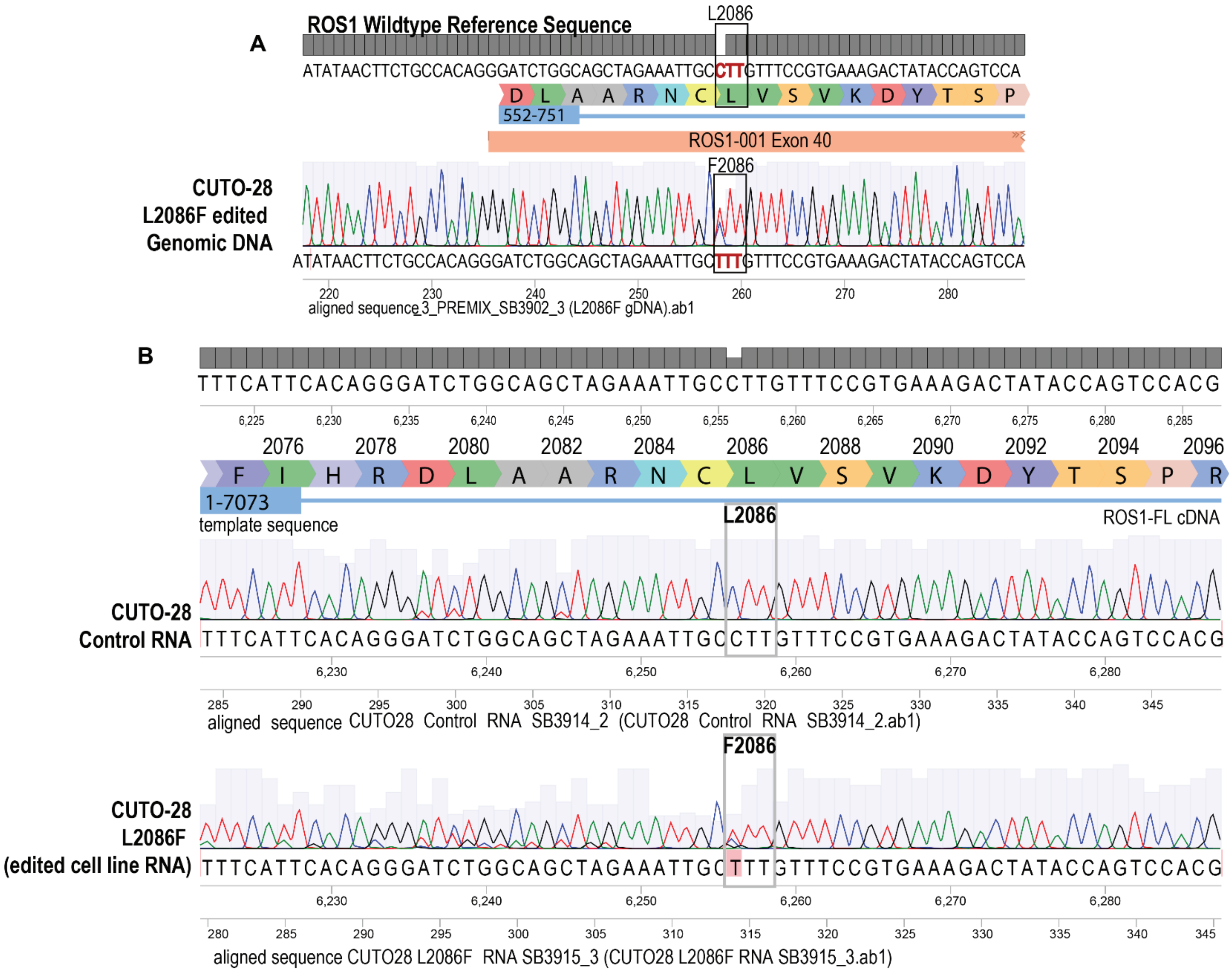

### Supplementary Figure 5

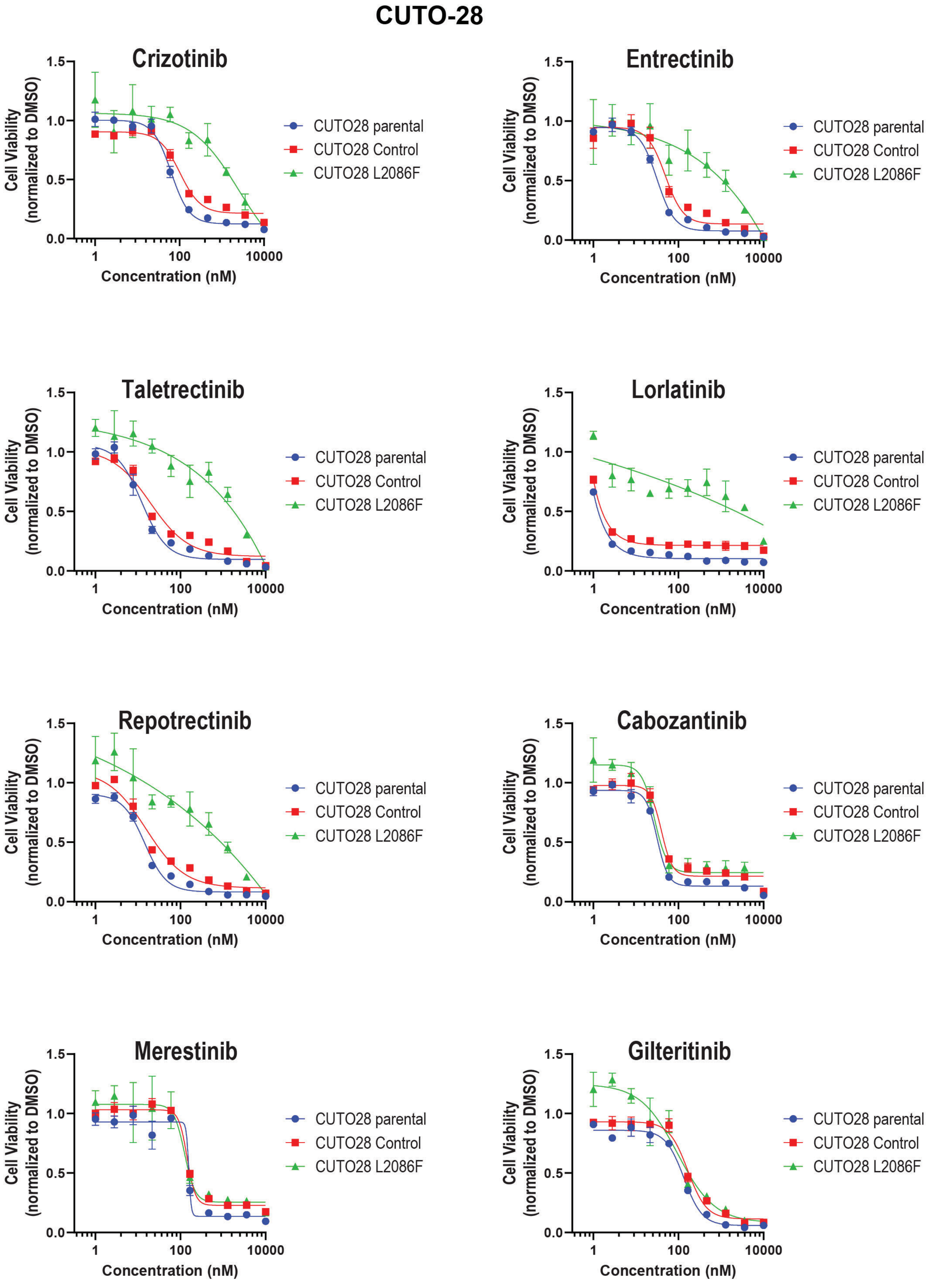

**SUPPLEMENTARY TABLE S1**

| Primer Name | Primer Sequence (all indicated 5' - 3') |
|---|---|
| ROS1 F2004C S | GGCACATCTGATGAGCAAATGTAATCATCCCAACATTCTG |
| ROS1 F2004C AS | CAGAATGTTGGGATGATTACATTTGCTCATCAGATGTGCC |
| ROS1 L2026M F | CCTCCATCAGTTCCATGATAATGTATTGGGGTTCATTCA |
| ROS1 L2026M R | TGAATGAACCCCAATACATTATCATGGAACTGATGGAAGG |
| ROS1 G2032R AS | AATAAGTAAGAAGGTCTCTTCCCTCCATCAGTTCCAG |
| ROS1 G2032R S | CTGGAACTGATGGAGGGAAGAGACCTTCTTACTTATT |
| L2086F_REV | AGTCTTTCACGGAAACAAAGCAATTTCTAGCTGCCAG |
| L2086F_FWD | CTGGCAGCTAGAAATTGCTTTGTTTCCGTGAAAGACT |
| L2086FguideF | caccgTATAGTCTTTCACGGAAACA |
| L2086FguideR | aaacTGTTTCCGTGAAAGACTATAc |
| L2086seqF | taatgcaggcagggcttaaa |
| L2086seqR | tgctgagacccaggaatagta |
| ROS1 F10 | AGAAGGGTTCCACAGACCAG |
| ROS1 R13 | TCCAGTCTCCCTCCTGTTTG |
| L2086F HDR template | ATTTCTGGAACTttattgttgttattattattattattaAATATAACTTCTGCCAC<br>AGGGATCTGGCAGCTAGA<br>AATTGCtTTGTTTCCGTGAAAGACTATACCAGTCCACGGATAGTGAA<br>GATTGGAGACTTTGGACTCG<br>CCAGAGACATCTATAAAAATGA |
